# Animal soil food web complexity triggers shifts in microbial communities, including PAH degraders, but without clear effects on phenanthrene phytoremediation

**DOI:** 10.1101/2021.12.14.472730

**Authors:** Sara Correa-Garcia, Vincenzo Corelli, Julien Tremblay, Jessica Ann Dozois, Eugenie Mukula, Armand Séguin, Etienne Yergeau

**Author notes:** Address correspondence to prof. Etienne Yergeau 531, boulevard des Prairies, Laval, QC, H7V 1B7, Canada.

## Abstract

The aim of this study was to determine whether the complexity of the animal soil food web (SFWC) is a significant factor influencing the soil microbial communities, the productivity of the willow, and the degradation rates of 100 mg kg^-1^ phenanthrene contamination. The SFWC treatment had eight levels: just the microbial community (BF), or the BF with nematodes (N), springtails (C), earthworms (E), CE, CN, EN, CEN. After eight weeks of growth, the height and biomass of willows were significantly affected by the SFWC, whereas the amount of phenanthrene degraded was not affected, reaching over 95% in all pots. SFWC affected the structure and the composition of the bacterial, archaeal and fungal communities, with significant effects of SFWC on the relative abundance of fungal genera such as *Sphaerosporella*, a known willow symbiont during phytoremediation, and bacterial phyla such as *Actinobacteriota*, containing many PAH degraders. These SFWC effects on microbial communities were not clearly reflected in the community structure and abundance of PAH degraders, even though some degraders related to the *Actinobacteriota* and the diversity of Gram-negative degraders were affected by the SFWC treatments. Overall, our results suggest that, under our experimental conditions, SFWC does not affect significantly willow phytoremediation outcomes.

**Importance:** Polycyclic aromatic hydrocarbons (PAH) pose a threat to soil ecosystems. Phytoremediation is a green technology that can help restore ecosystems’ health affected by PAH contamination. Past research on phytoremediation of PAH has focused on the roles of plant and microbes in contaminant fates. However, soil environments usually harbor large faunal communities that interact with both the plant and the microbial communities, potentially altering the phytoremediation process. We hypothesized that soil food web complexity (SFWC), represented by increasing levels of soil fauna, influences PAH degradation through interactions with the plant and the microbes. In this study, we demonstrated that SFWC increases the plant biomass and changes the composition of the microbial community, especially the fungal structure. In our pot experiment, more complex levels of SFWC did not contribute to higher degradation rates, but the increased plant biomass and the higher relative abundance of fungal genera associated with lower plant stress could indirectly contribute to better phytoremediation outcomes. Our results highlight the importance of considering the contributions of soil fauna to the success of phytoremediation.

## 1. Introduction

Polycyclic aromatic hydrocarbons (PAH) are organic compounds originating from the incomplete combustion of organic matter. They are considered as contaminants in the environment (Jones et al., 1989; Ravindra et al., 2008), especially in soils (Wilcke, 2007), due to their detrimental effects on ecosystems (Menzie et al., 1992). Phenanthrene (PHE) is a PAH formed by three fused benzenic rings. It has been widely used as a model molecule, alone or with other PAHs in degradation experiments (Erktan et al., 2020b; Li et al., 2019; Lu et al., 2019; Sun et al., 2010).

Phytoremediation has demonstrated a significant potential to tackle PAH contamination (Guo et al., 2018; Kuppusamy et al., 2017; Lu and Lu, 2015; Pilon-Smits, 2005; Spriggs et al., 2005). Rhizoremediation, the process where microorganisms are stimulated to degrade contaminants in the rhizosphere environment by root exudates (Correa-García et al., 2018), is one of the main phytoremediation approaches used to degrade PAH. Different species from the *Salicaceae* family have been used for the phytoremediation of hydrocarbons in the Canadian climatic context (Gonzalez et al., 2018; Tardif et al., 2016; Yergeau et al., 2018, 2015, 2014), including PHE (Correa-García et al., 2021). Particularly, willow trees develop deep root systems, present rapid growth rates and are resistant both to biotic and abiotic stressors. These characteristics make them suitable for rhizoremediation.

Many soil microorganisms metabolize PHE and other PAH (Cerniglia, 1993). Under aerobic conditions, PHE is catabolized into catechols and further TCA cycle intermediates (Cerniglia, 1993; Eaton and Chapman, 1992; Mallick et al., 2011). This degradation process starts with the incorporation of a molecular oxygen into one of PHE aromatic nucleus by a ring hydroxylating dioxygenase (RHD) multicomponent enzyme system (Kauppi et al., 1998). The genes coding for the alpha subunit of the PAH-RHD form a monophyletic group (Habe and Omori, 2003). Hence, these genes are often used to monitor bacterial degrader communities in PAH contaminated environments using PCR primers designed for either Gram-negative or Gram-positive bacteria (Cébron et al., 2008). Many bacteria contain RHD genes are commonly found in the root systems of *Salicaceae* trees (Bell et al., 2015, 2014a; Correa-García et al., 2021; Khan et al., 2014; Pagé et al., 2015). However, the outcome of phytoremediation often remains unpredictable and variable (Bell et al., 2014b; Correa-García et al., 2018). This variability is partially due to the initial soil microbial community composition and diversity (Bell et al., 2015, 2013; Yergeau et al., 2015) and soil physicochemical characteristics (Correa-García et al., 2021).

Interestingly, both factors can be affected by soil animal communities. For instance, collembolans and nematodes feed on soil bacteria, fungi and secondary roots, which can stimulate the growth of specific microbial taxa (Crowther et al., 2011; Endlweber et al., 2009; Erktan et al., 2020b, 2020a). Collembolans and nematodes were also shown to stimulate the growth of secondary plant roots mainly through changes in the availability of nutrients (Alphei et al., 1996; Endlweber and Scheu, 2006; Ngosong et al., 2014). Earthworms, as ecosystem engineers, have a significant impact on soil physicochemical characteristics, due to their role in organic matter degradation, nutrient turnover, soil oxygenation and microbial movement between the bulk soil and the rhizosphere (Bartlett et al., 2010; Lavelle and Spain, 2001; Spain et al., 1992; Yang and van Elsas, 2018).

Consequently, collembolans, nematodes, earthworms and the rest of the soil fauna may have an interesting, yet unexplored, role to play in the rhizoremediation of PAH (Li et al., 2015; Zeb et al., 2020), probably through changes in soil microbial communities and/or soil physicochemical properties. Still, just a few studies have explore the role of soil fauna during remediation of PAH, and the majority of the experiments have only considered single species or single ecological groups (Jing et al., 2017; Zhou et al., 2019). However, many animal groups survive in contaminated soil matrices (Delgado-Balbuena et al., 2016; Jing et al., 2017; Sun et al., 2017; Zavala-Cruz et al., 2013). Despite these hints, little is known about the possible effect of trophic complexity of soil food webs in the fate of PAH in soils. In view of their essential roles in soil ecosystems, soil animals and the complexity of their food web interactions could also affect the outcome of phytoremediation.

Here, we hypothesized that more complex soil food webs result in more efficient phytoremediation, through specific changes in the rhizosphere microbial community structure. Specifically, we hypothesized that more complex food webs under phenanthrene contamination would result in 1) larger plants, 2) larger shifts in microbial communities when compared to non-contaminated controls and, consequently, 3) more degradation. Therefore, we aimed at assessing the response of microbial communities and willow trees to increasingly complex soil animal food webs during PHE rhizoremediation. We performed a PHE rhizoremediation pot experiment consisting of a full factorial combination of collembolans, nematodes and earthworms presence/absence. After eight weeks of willow growth, the 16S rRNA gene of Bacteria and Archaea, the ITS1 region of Fungi and the PAH-RHDα genes of Gram-negative and Gram-positive bacteria were sequenced, the absolute abundance of PAH-RHDα genes was measured and the willow height and biomass and the residual concentration of phenanthrene were determined.

## 2. Material and Methods

### 2.1. Soil and biological material

The soil used was acquired from Savaria Matériaux paysagers Ltée (Laval, QC, Canada) and it was subjected to gamma irradiation at a dose ranging from 12.4 to 24.8 kGy by Nordion (Laval, QC, Canada) to significantly reduce the community of soil dwelling invertebrates whilst preserving a significant fraction of the microbial diversity and abundance (McNamara et al., 2003). The irradiated soil was subjected to funnel extractions with heat irradiation to confirm the effectiveness of the gamma irradiation. No arthropods were recovered after 48h of extraction, thus confirming the almost complete reduction of living soil invertebrates’ communities. A wet extraction to recover nematodes was not performed since previous experiments report recovering 100% empty carcases of nematodes several weeks after soil irradiation with doses as low as 3 kGy (Buchan et al., 2012; Gebremikael et al., 2015). The control potting mix was prepared by mixing perlite and the irradiated soil in a proportion of 1:2 (v:v) to improve permeability and prevent soil compaction. For the contaminated soil, after gamma irradiation, the soil was spiked with 100 mg·kg^-1^ dry soil of phenanthrene according to the following protocol. Batches of 1 kg of dried, 2 mm sieved soil were spiked with 1 g of phenanthrene (Sigma Aldrich) diluted in 100 mL of acetone. Batches of control soil were also spiked with acetone alone. The spiked soil was left in a chemical hood for 48h until the acetone was completely evaporated. Then, the spiked 1 kg batches (contaminated or control) were thoroughly incorporated into 9 kg of uncontaminated soil in a cement mixer. After that, perlite was mixed into the soil in a proportion of 1:2 (v:v) until the potting mix reached homogeneity.

Willow cuttings (*Salix purpurea* cv. FishCreek) were acquired from Agro Énergie (Saint-Roch-de-l’Achigan, Quebec, Canada). Prior to planting, willow cuttings were pre-soaked by submerging 80% of their length in tap water at room temperature for one week. Cuttings showing signs of disruption of dormancy (greener buds, incipient root tips) were selected for the experiment.

*Aporrectodea caliginosa* adult earthworm specimens with a fully developed clitellum were collected in April 2018 on the INRS campus and were kept in soil microcosms composed of 20×20×12 cm plastic boxes containing 500 g of 2 mm sieved soil with a pierced cover. Up to 10 adult earthworms were kept in each microcosm. This system was enriched with 100 g of cattle manure and the soil humidity was kept at 30% by weighing the boxes and adding the corresponding amount of tap water every two weeks. Five adults presenting a fully developed clitellum were deposited at the surface of the corresponding pots.

Adult nematodes from *Caenorhabditis sp*. were isolated from a decomposing earthworm in our soil microcosms. Isolation in agar culture plates, identification and culture conditions for the nematodes were carried out as described in (Barrière and Félix, 2006). Briefly, the decomposing earthworm body where the nematodes were found was placed in a standard Nematode Growth Medium (NGM) agar Petri dish containing an *E. coli* OP50 lawn (Brenner, 1974). The Petri dish was moistened with sterile water to facilitate migration. Up to 30 individual worms were collected from the bacterial lawn after times ranging from 15 min to 2 h and placed with a worm picker into a new individual agar plate. The Petri dishes were kept at room temperature. After 7 days, *Caenorhabditis sp*. worms were identified under the dissecting microscope discriminating the different families by their oral apparatus, the color of the intestinal cell contents (light brown) and the central position of the vulva. Cultures were maintained by transferring nematodes every 2-4 days into a fresh agar plate for 2 weeks prior to the beginning of the experiment. A water solution of nematodes was prepared and 5 mL of it, containing approximately 5,000 individuals of all ages, were inoculated into the soil. This amount was calculated by mounting 10 µL of the solution in a slide and counting individuals under the stereoscope. A total of 4 slides were mounted and used to calculate the mean. The number of observed individuals ranged between 8 and 12 per slide. Springtails from *Tomocerus sp.* and *Folsomia candida* were used as the collembolans representatives. *Tomocerus* culture were purchased at Magazoo (Montréal, QC, Canada).

*Folsomia* springtails were extracted from the same soil used in this experiment through heat irradiation with a funnel and kept in culture conditions at room temperature for 2 weeks prior to the beginning of the experiment. The springtail cultures consisted of coconut fiber and charcoal moistened once a week in 500 mL tubular plastic containers with a pierced lid covered with light filter paper to allow air exchange. Before inoculation, springtail cultures were placed at 4 °C overnight to reduce their metabolic rate and allow handling of the colonies (Cooper, 2011). The colonies were mixed, including the substrate, and were subsequently divided in 55 individual bags of approximately 100 g. Springtails of 4 bags were extracted (funnel extraction with a clamped rubber tube for 24 h under incandescent lamp) and counted, showing that each bag contained approximately 200 to 250 individuals. To keep the springtails anesthetised, the bags were kept at 4 °C in ice until inoculation where one bag per pot was used.

### 2.2. Experimental design

A full factorial experiment consisting of three factors was implemented: contamination, food web complexity and plant compartment. The contamination treatment consisted of two levels: soils contaminated with 100 mg·kg^-1^ dry soil of phenanthrene (PHE) and an uncontaminated control soil (CTRL). The soil was gamma-irradiated to allow for the control of the food web complexity treatment. The food web complexity treatments consisted of eight levels: bacteria and fungi as control (BF, the irradiated soil only), the irradiated soil (BF) plus springtails (C), nematodes (N) or earthworms (E), and BF plus all the possible two and three animal combinations (CE, CN, EN, CEN). Microbial communities and phenanthrene concentrations were evaluated in two soil compartments: bulk (Bulk) and rhizosphere (Rhizo) soils. The two first factors (contamination and food web complexity) resulted in 16 soil treatments that were used for the pot experiment. The pots were arranged in six experimental blocks, wherein the 16 treatments were randomly distributed, for a total of 96 pots that were placed outside at the Centre Armand Frappier Santé Biotechnologie (INRS, Laval, QC, Canada, 45.541393°N, - 73.716980°W). One cutting was planted per six-liter pot containing approximately 4 kg of potting mix on the 7^th^ and 8^th^ of August 2018. Animals were added at the surface of the potting mix (N and E) or at 5 cm under the surface (C), immediately after the cuttings were planted. Pots were then covered with coconut fiber to prevent animals from escaping and to reduce phenanthrene evaporation and photooxidation. Then, the pots were connected to an automated drip irrigation system and received approximately 400 mL of water every day at 8 am.

### 2.3. Sampling and plant trait measurements

The experiment was sampled between the 1^st^ and 4^th^ of October 2018, after approximately 8 weeks of plant growth. Plant height and the number of shoots were measured prior to clipping aboveground shoot biomass (excluding the original cutting). Afterwards, the cuttings with the attached roots were taken out from the pots and rhizosphere and bulk soil samples were collected for microbial community analysis and phenanthrene concentration measurements as described in Correa-García et al., (2021), resulting in 192 soil samples. Soil samples were kept at 4 °C until transportation to the lab (around 2 h) where they were placed at −20 °C. For all plants harvested, aboveground willow biomass was weighed fresh, oven-dried for 24 h at 60 °C and weighed again to recover dried biomass and aboveground water content values. Root biomass was collected sieving at 2 mm all the soil from every pot to ensure recovery of the maximum amount of root biomass. Roots collected were subsequently cleaned from perlite and adhered soil, then dried and weighed. The objective of our study was not to determine the effects of PHE on the survival and fitness of soil invertebrates, therefore no animal extraction was performed.

However, the presence of variable numbers of surviving E and C was visually confirmed on the pots receiving these treatments at the end of the experiment.

### 2.4. Phenanthrene quantification

Phenanthrene concentration was measured in the rhizosphere and bulk soil samples taken at the end of experiment. Two phenanthrene measurements were taken per biological replicate. The phenanthrene extraction protocol consisted on an in-house method developed in the lab and described in Correa-Garcia (2021). Briefly, 4 g of frozen soil were mixed with 900 µL of ethyl acetate, 3 mL of distilled water and 10 ppm of phenanthrene-d_10_ in 100 µL of ethyl acetate used as internal standard. Then, the samples were placed in an ultrasonic bath at 60 kHz for 15 min. Next, the samples were shaken overnight at 300 rpm at room temperature. Afterward, the samples were centrifuged at 270 g for 10 min and the organic phase was recovered. The organic phase was further purified by centrifugation at 5,000 × g for 1 min to yield the final phenanthrene extracts to be used in GC-MS analysis.

Phenanthrene extracts were analyzed using a Trace GC Ultra system (Thermo Scientific) with a 30 x 0.25 mm (0.25 µm thickness) DB-5 MS capillary column (Agilent J & W capillary GC) and coupled to a Polaris Q benchtop Ion Trap Mass Spectrometer. The injector temperature was set at 250°C and the analyzer at 350°C. The GC-MS program consisted in 2 min hold at 70°C, increasing temperature to 310°C at 30°C min^-1^ followed by 6 min hold at 310°C. Helium was used as carrier gas at a flow rate of 0,3 mL/min. The injection volume was 3 µL. MS scan range was set at 70-600 m/z. Standard calibration curves for serial concentrations ranging from 1 ppm to 100 ppm were calculated by plotting the peak areas against the concentration of reference. The final concentration of phenanthrene is expressed as mg kg^-1^.

### 2.5. Microbial community composition: DNA isolation and library preparation

The DNA isolation steps and the library preparation follow the steps described in-depth in Correa-García et al., (2021). Briefly, 250 mg per sample of bulk or rhizosphere soil were transferred to a 2 mL Powerbead Pro tube. The samples were then homogenized in a FastPrep®- 24 (MP Biomedicals) at 6 m/s for two times 45 s. Then, DNA was extracted using the DNeasy® Powersoil® kit (Qiagen) following the protocol. The V3-V4 regions of the 16S rRNA gene were amplified using the primer set 515F – 806R (Caporaso et al., 2012) for bacterial and archaeal communities. The internal transcribed spacer region (ITS) with the primer set ITS1F – 58A2R (Martin and Rygiewicz, 2005) were used to target the fungal community. The PAH-RHDα gene clusters were amplified using the primer set 610F – 916R for Gram negative (GN) bacteria and using the primer set 641F – 933R for Gram positive (GP) bacteria (Cébron et al. 2008). All microbial regions were amplified in 25 µL volumes containing 10 - 20 ng of DNA template as in Correa-García et al., (2021). The PCR conditions for bacteria and archaea were as follows: initial denaturation at 95°C for 5 min; 35 cycles at 95°C for 30 s, 55°C for 30 s, 72°C for 1 min, and a final elongation phase at 72°C for 5 minutes. For fungi, the annealing temperature was set at 59°C instead of 55°C, and the number of cycles at 30 instead of 35. PAH-RHDα genes had annealing temperatures set at 57°C and 54°C for GN and GP primer sets, respectively and a total of 30 cycles. PCR products were cleaned following the Illumina’s protocol “16S Metagenomic Sequencing Library preparation” guide (Part #15044223 Rev. B). Then, clean PCR products were tagged using 400 nM of each Nextera XT unique index primers under the following thermal cycling conditions: 95°C initial denaturation phase for 5 min, followed by 8 cycles consisting of 95°C denaturation for 30 s, annealing at 55°C for 30 s, elongation at 68°C for 30 s, and a final elongation phase at 68°C during 5 min. The clean amplicons were pooled in equimolar ratios. These libraries were sequenced on an Illumina Miseq sequencer at the Centre d’expertise et de services Génome Québec (Montréal, QC, Canada).

### 2.6. Real-time PCR quantification of PAH degrading genes

The real-time quantitative PCR (qPCR) was conducted on a Stratagene Mx3005P qPCR system (Agilent Technologies). The qPCR reactions were performed using the primers designed by Cébron et al. (2008) as in Correa-García et al., (2021). Briefly, the denaturation step at 95°C for 5 min was followed by 40 cycles of denaturation at 95°C for 30 s, annealing for 35 s at either 57°C (GN) or 54°C (GP) and an elongation step at 72°C for 75 s, after which the SYBR Green signal intensity was measured. A melting curve analysis was performed where signal intensity was measured at 0.5°C increments every 5 s from 51 to 95°C. Standards were made from 10-fold dilutions of linearized plasmid containing the gene fragment of interest, cloned from amplified from soil DNA (Yergeau et al. 2009). At the end of the run, the Ct values were evaluated, and the gene copy numbers were calculated from the standard curve based on the Ct values. The efficacy of the qPCRs ranged from 49.6% (R^2^ = 0.994) to 85.0% (R^2^ = 0.996).

### 2.7. Bioinformatic analyses

Sequences were analyzed using AmpliconTagger (Tremblay et Yergeau, 2019). Briefly, raw reads were scanned for sequencing adapters and PhiX spike-in sequences. Remaining reads with an average quality (Phred) score lower than 20 were discarded. The rest of the sequences were processed for generating Amplicon Sequence Variants (ASV; DADA2 v1.12.1; Callahan *et al*. 2016). Chimeras were removed with DADA2’s internal removeBimeraDeNovo(method=”consensus”) method followed by UCHIME reference (Edgar et al., 2011). Only ASVs with an abundance across all samples higher than 5 were retained. A global read count summary throughout the pipeline is provided in Suppl. Table 1 for all datasets.

ASVs were assigned a taxonomic lineage with the RDP classifier (Wang et al., 2007) using an in-house training set based on the complete Silva release 138 database (Quast et al., 2013) supplemented with eukaryotic sequences from the Silva database and a customized set of mitochondria, plasmid and bacterial 16S sequences. For ITS ASVs, a training set containing the Unite DB to classify sequences (sh_general_release_s_04.02.2020 version) was applied. The final lineages were reconstructed using the taxonomic depths having a score ≥ 0.5 (from a 0 to 1 range assigned by the RDP classifier). Taxonomic lineages were combined with the cluster abundance matrix obtained above to generate a raw ASV table, from which a bacterial/fungal organisms ASV table was generated.

PAH-RHDα GP and GN amplicon sequencing libraries were processed as described above up to the quality filtering step and remaining sequences were processed to generate ASVs (DADA2 v1.12.1) (Callahan et al., 2016). Custom RDP classifier training sets were generated for both PAH-RHDα GP and GN amplicon data types as follow. Each ASV was blasted (BLASTn) against the NCBI nt database (downloaded on February 21, 2020) with –max_target_seqs set to 20. Blast output was filtered to keep hits using the following thresholds: an e-value <= 1e-20, alignment length of at least 100 bp and alignment percentage of at least 60 %. Taxonomic lineages of each filtered blast hit were fetched from the NCBI taxonomy database. RDP training sets were generated as previously described (https://github.com/jtremblay/RDP-training-sets).

### 2.8. Statistical analyses

Statistical analyses were performed in R (v.3.5.0) (R Core Team, 2021) with the stats package (R Core Team, 2021). The differences in plant trait results, qPCR quantifications and phenanthrene concentration were assessed using ANOVA followed by *post hoc* Tukey HSD tests. Kruskal-Wallis analysis of variance coupled with the Dunn test as *post hoc* when the assumptions of normality and homoscedasticity were not met. The ANOVA test with White’s correction for heteroscedastic data was used to test differences in the relative abundance of genera with the car package (Fox and Weisberg, 2019).

Shannon H’ diversity index and species richness (observed number of ASV) were calculated with the otuSummary package (Yang, 2020) and tested with ANOVA.

The differences in the community structure were visually assessed using principal coordinate analyses (PCoA) with normalized ASV tables and the Bray-Curtis dissimilarity index calculated with the vegan package (Oksanen et al., 2019). The effect of the treatments on the microbial community structures was tested using PERMANOVA analyses with 999 permutations using the Adonis function. All graphs were created with the ggplot2 package (Wickham et al., 2019).

### 2.9. Data deposition

Raw sequencing reads were deposited in the NCBI SRA under BioProject accession PRJNA700608.

## 3. Results

### 3.1. Plant biomass and phenanthrene concentration

The willow shoot fresh biomass was mainly influenced by contamination (*F* = 17.806, *p* < 0.001), with CTRL presenting significantly higher biomass (from 50.542 g in CEN to 37.773 g in N) than PHE pots (44.510 g in CEN to 24.167 g in BF; Table 1). Generally, the willows with the highest biomass were growing in the CTRL soil, apart from the CEN PHE willows, which had the third highest biomass of all treatments (Table 1). The soil food web complexity (SFWC) was also a significant factor (*F* = 3.531, *p* = 0.002), with higher biomass values in the CEN treatment for both contamination levels.

**Table 1.**
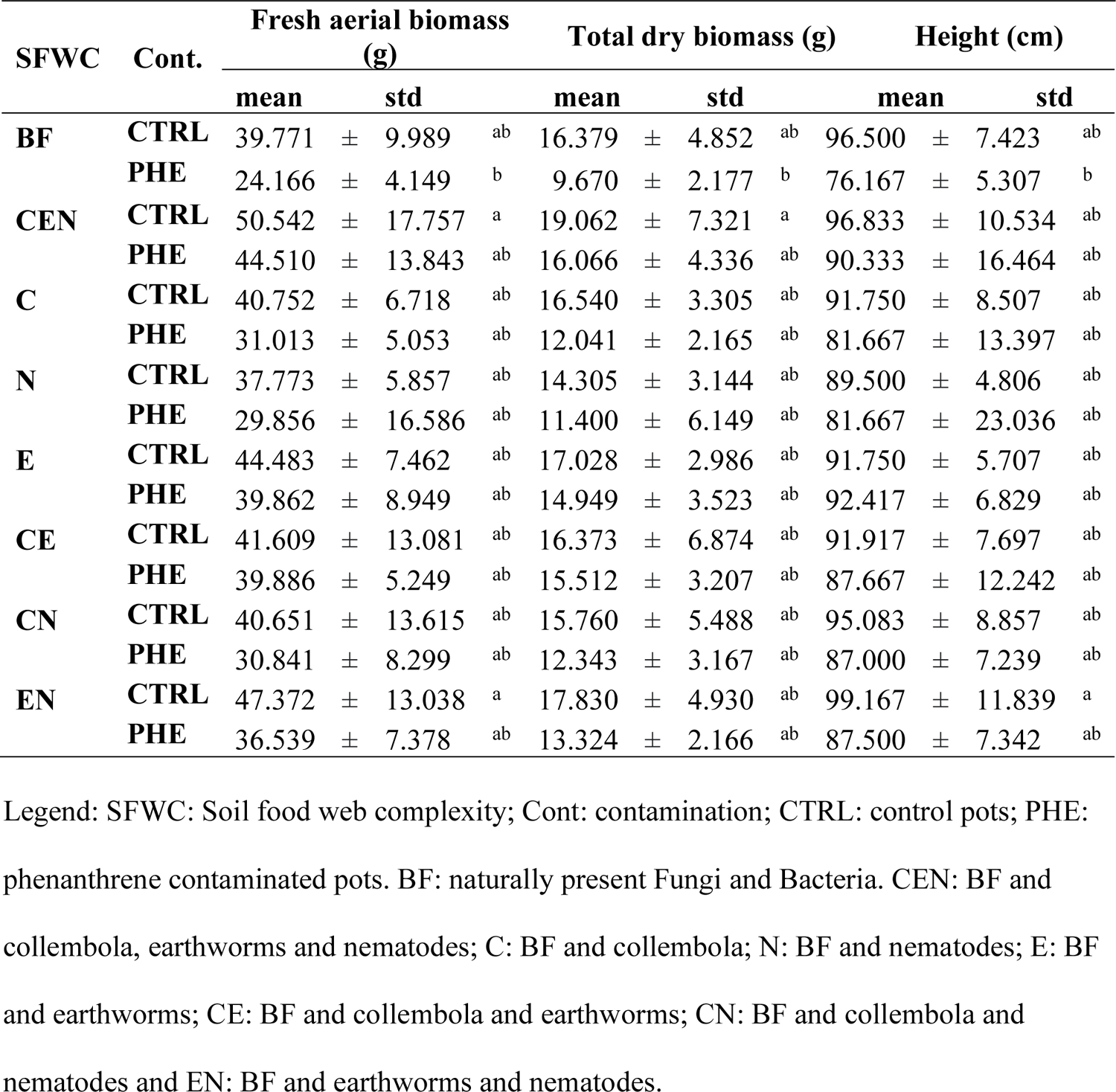
Mean and standard deviation for willow morphological traits. Letters denote significantly different groups as stated by the Tukey post hoc test.

Willow total dry biomass was significantly higher in CTRL (from 19.062 g in CEN to 14.305 g in N) than in PHE soils (16.066 g in CEN to 6.670 g in BF) (*F* = 19.725, *p* < 0.001; Table 1). SFWC also had a significant effect on willow total dry biomass (*F* = 2.147, *p* = .048). Again, the PHE CEN willows had similar biomass to willows growing in CTRL soil, and all willows growing in PHE soils with earthworms had higher total biomass than the willows growing in PHE soils without earthworms (Table 1). Only the total biomass of the CRTL CEN and the PHE BF treatments were significantly different in *post-hoc* tests.

Contamination was the only factor significantly affecting the willow height (*F* = 15.176, *p* < 0.001). Again, in general, CTRL plants (from 99.167 cm in EN to 89.500 cm in N) were higher than PHE plants (from 92.417 cm in E to 76.167 cm in BF), but willows growing in PHE soil in the presence of earthworms were taller than in the absence of earthworms (Table 1). For the three plant parameters, only BF PHE was significantly lower than the best performers of the CTRL groups (Table 1).

There were significant differences in terms of phenanthrene degradation between compartments (*F* = 8.721, *p* = 0.004), with slightly more phenanthrene degraded in the bulk soil. There was also a nearly significant effect of the compartment x SFWC interaction term (*F* = 1.848, *p* = 0.091). However, these differences were not biologically significant, since the highest phenanthrene value retrieved was of 3.99 mg kg^-1^ and the lowest 0.89 mg kg^-1^ (Figure 1), representing overall degradation rates ranging between 96 and 99% of the 100 mg kg^-1^ phenanthrene applied, both in the bulk and rhizosphere soils.

**Figure 1.**
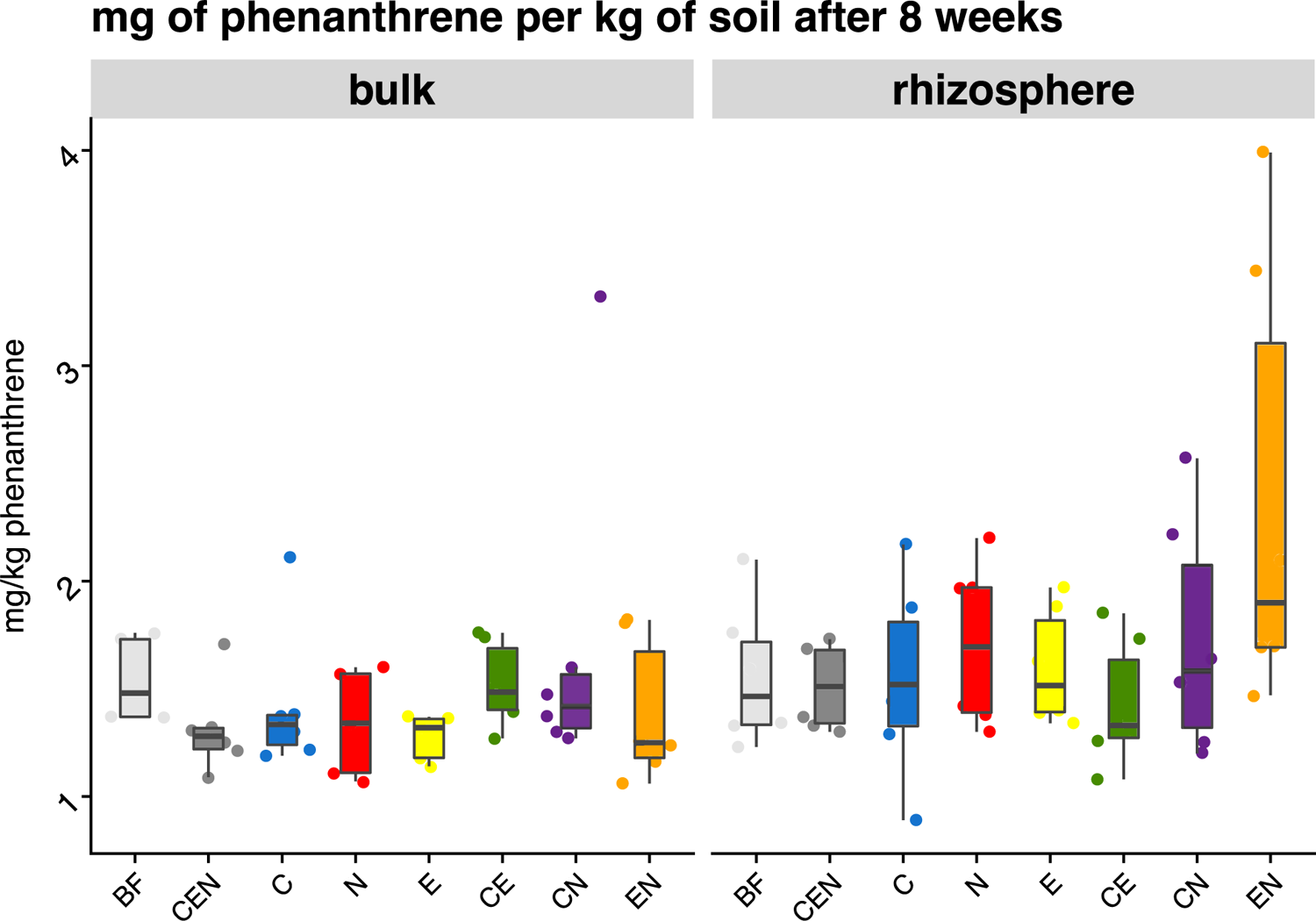
Quantification of phenanthrene (as mg kg ^-1^ soil) after 8 weeks of willow growth in bulk and rhizosphere soil compartments for all soil food web complexity (SFWC) treatments. No significant differences were found (n = 6). Legend: SFWC levels: BF = only naturally present *Bacteria* and *Fungi*; CEN = BF plus added collembola, nematodes and earthworms; C = BF and added collembola; N = BF and added nematodes; E = BF and added earthworms; CE = BF and added collembola and earthworms; CN = BF and added collembola and nematodes; EN = BF and added nematodes and earthworms.

### 3.2. Microbial community diversity

The median values for the Shannon H’ diversity index and the number of observed ASV for all treatments and soil compartments are presented in Table 2 and the statistical analysis on the observed number of ASV are presented in the Suppl. Table 2. The contamination and the soil compartment significantly affected the fungal Shannon H’ diversity, but the SFWC treatment also had a minor effect (Table 3). The mean Shannon H’ diversity was the highest in N (1.79) and CEN (1.66) soils, whilst the lowest was CN (0.96). In general, PHE and bulk soils had lower fungal Shannon H’ diversity.

**Table 2.** Median Shannon H’ diversity and median ASV. Legend as described in Table 1.

|  | ITS<br>gene<br>(n=6) |  | 16S rRNA<br>gene<br>(n=6) |  | PAH - RHD Gram<br>Negative gene<br>(n=6) |  | PAH - RHD Gram<br>Positive gene<br>(n=6) |  |
| --- | --- | --- | --- | --- | --- | --- | --- | --- |
|  | Shannon | ASV n° | Shannon | ASV n° | Shannon | ASV n° | Shannon | ASV n° |
| **BF bulk CTRL** | 1.480 | 75 | 5.045 | 463 | 2.766 | 29.5 | 4.944 | 227 |
| **BF bulk PHE** | 0.746 | 53.5 | 5.446 | 473.5 | 3.466 | 62.5 | 5.127 | 242 |
| **BF rhizosphere CTRL** | 1.617 | 108 | 5.350 | 535 | 3.496 | 51 | 5.108 | 281.5 |
| **BF rhizosphere PHE** | 1.643 | 136.5 | 5.215 | 498 | 3.360 | 40.5 | 5.572 | 409.5 |
| **CEN bulk CTRL** | 2.102 | 82 | 4.920 | 337 | 2.581 | 20 | 4.854 | 189.5 |
| **CEN bulk PHE** | 0.944 | 50 | 5.411 | 473 | 3.580 | 52.5 | 5.197 | 225 |
| **CEN rhizosphere CTRL** | 2.835 | 153 | 5.175 | 482 | 3.050 | 35.5 | 5.035 | 218.5 |
| **CEN rhizosphere PHE** | 1.435 | 92 | 5.304 | 557 | 3.499 | 49.5 | 5.607 | 410 |
| **C bulk CTRL** | 1.233 | 61.5 | 4.723 | 342.5 | 2.675 | 17.5 | 4.674 | 137 |
| **C bulk PHE** | 0.723 | 34.5 | 5.118 | 371.5 | 3.288 | 46 | 5.017 | 209 |
| **C rhizosphere CTRL** | 1.255 | 97 | 5.189 | 455.5 | 3.267 | 37 | 5.242 | 281 |
| **C rhizosphere PHE** | 1.101 | 87 | 5.312 | 477.5 | 3.479 | 50 | 5.585 | 409.5 |
| **N bulk CTRL** | 2.428 | 91 | 5.037 | 377 | 3.276 | 42 | 4.758 | 164 |
| **N bulk PHE** | 0.555 | 29.5 | 5.391 | 502 | 3.121 | 38.5 | 5.158 | 249 |
| **N rhizosphere CTRL** | 1.594 | 162 | 5.211 | 495.5 | 2.727 | 20 | 5.330 | 301.5 |
| **N rhizosphere PHE** | 1.557 | 153.5 | 5.463 | 533.5 | 3.484 | 54.5 | 5.633 | 436 |
| **E bulk CTRL** | 1.056 | 75.5 | 5.207 | 418.5 | 3.152 | 28.5 | 4.781 | 158 |
| **E bulk PHE** | 0.669 | 39.5 | 5.330 | 477 | 3.447 | 52 | 5.163 | 230 |
| **E rhizosphere CTRL** | 2.410 | 120 | 5.222 | 522 | 2.588 | 22.5 | 4.933 | 185 |
| **E rhizosphere PHE** | 0.793 | 105 | 5.338 | 481 | 3.387 | 41 | 5.519 | 360 |
| **CE bulk CTRL** | 1.170 | 63 | 4.990 | 405.5 | 3.037 | 25 | 4.966 | 211.5 |
| **CE bulk PHE** | 0.835 | 33 | 5.311 | 429 | 3.063 | 31.5 | 5.479 | 328 |
| **CE rhizosphere CTRL** | 2.093 | 104 | 5.258 | 472.5 | 2.880 | 27.5 | 5.038 | 263 |
| **CE rhizosphere PHE** | 1.262 | 108 | 5.428 | 552 | 3.472 | 55 | 5.584 | 432.5 |
| **CN bulk CTRL** | 0.828 | 40 | 4.910 | 337 | 3.025 | 47.5 | 4.692 | 149.5 |
| **CN bulk PHE** | 0.658 | 28.5 | 4.914 | 315.5 | 3.178 | 32.5 | 5.233 | 243.5 |
| **CN rhizosphere CTRL** | 1.225 | 113.5 | 5.350 | 462 | 2.911 | 33.5 | 5.370 | 355 |
| **CN rhizosphere PHE** | 0.843 | 98.5 | 5.619 | 571.5 | 3.394 | 44 | 5.680 | 457 |
| **EN bulk CTRL** | 0.949 | 49 | 5.221 | 461 | 3.376 | 53 | 4.811 | 162.5 |
| **EN bulk PHE** | 0.530 | 25.5 | 5.323 | 458 | 3.404 | 55.5 | 5.135 | 209 |
| **EN rhizosphere CTRL** | 0.894 | 79 | 5.225 | 523.5 | 2.896 | 26.5 | 5.051 | 221 |
| **EN rhizosphere PHE** | 0.841 | 98 | 5.533 | 574.5 | 3.408 | 48 | 5.659 | 430 |

**Table 3.**
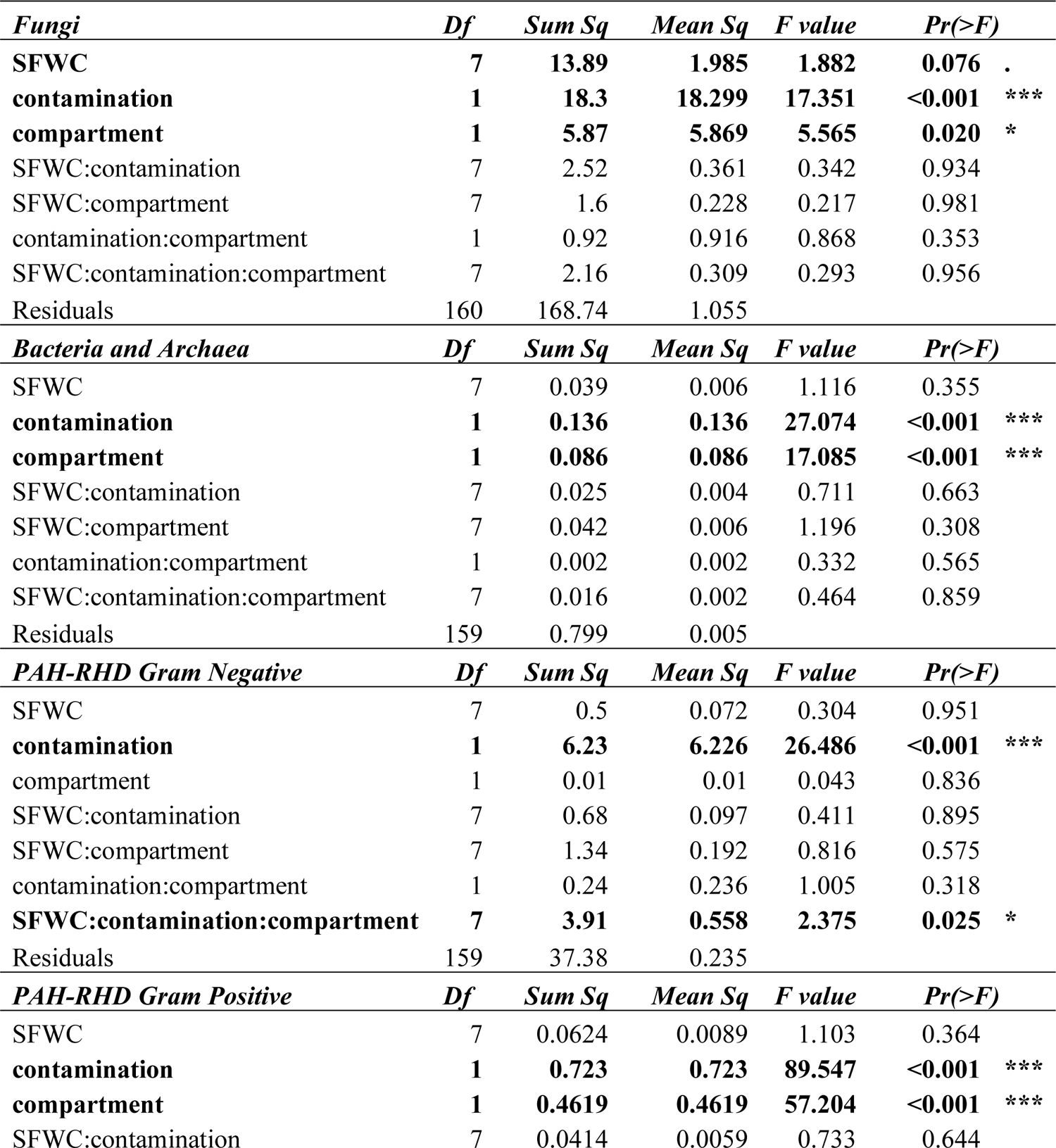
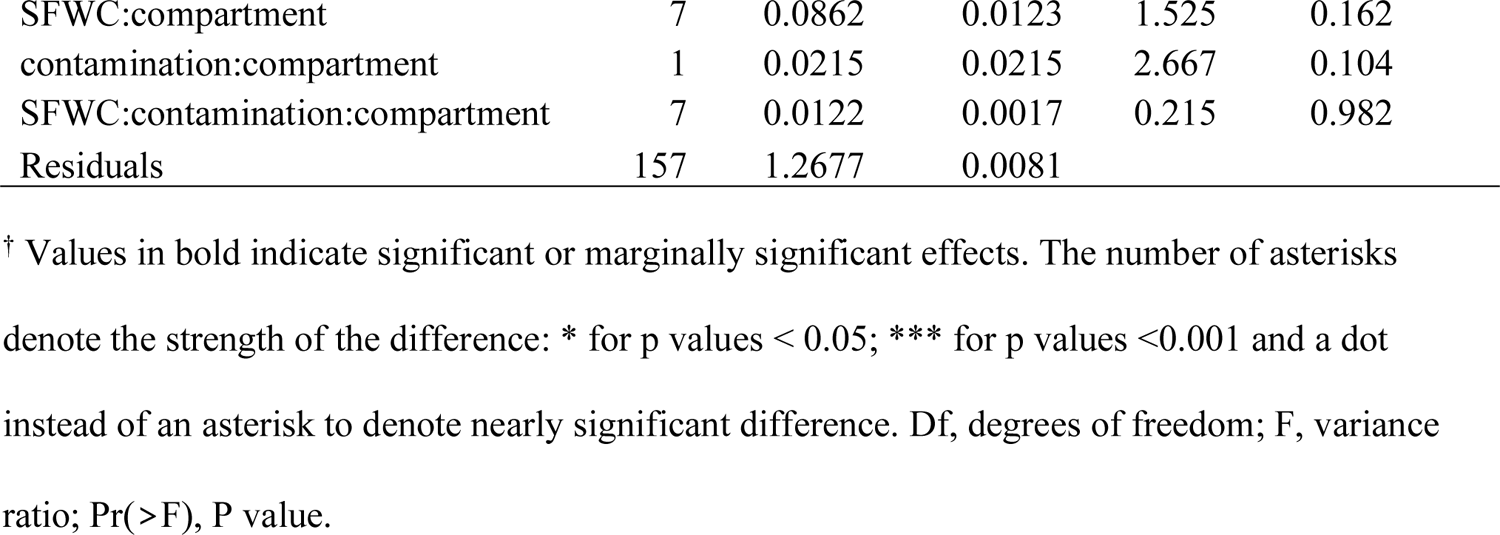
Summary of the three-way analysis of the variance (ANOVA) on the Shannon H’ diversity based on the ASVs of the ITS region for fungi, the 16S rRNA gene for total bacterial and archaea, and ASVs for Gram-negative and Gram-positive PAH-RHDα genes for bacterial degraders^†^.

| *Fungi* | *Df* | *Sum Sq* | *Mean Sq* | *F value* | *Pr(>F)* |  |
| --- | --- | --- | --- | --- | --- | --- |
| **SFWC** | **7** | **13.89** | **1.985** | **1.882** | **0.076** | . |
| **contamination** | **1** | **18.3** | **18.299** | **17.351** | **<0.001** | *** |
| **compartment** | **1** | **5.87** | **5.869** | **5.565** | **0.020** | * |
| SFWC:contamination | 7 | 2.52 | 0.361 | 0.342 | 0.934 |  |
| SFWC:compartment | 7 | 1.6 | 0.228 | 0.217 | 0.981 |  |
| contamination:compartment | 1 | 0.92 | 0.916 | 0.868 | 0.353 |  |
| SFWC:contamination:compartment | 7 | 2.16 | 0.309 | 0.293 | 0.956 |  |
| Residuals | 160 | 168.74 | 1.055 |  |  |  |
| *Bacteria and Archaea* | *Df* | *Sum Sq* | *Mean Sq* | *F value* | *Pr(>F)* |  |
| SFWC | 7 | 0.039 | 0.006 | 1.116 | 0.355 |  |
| **contamination** | **1** | **0.136** | **0.136** | **27.074** | **<0.001** | *** |
| **compartment** | **1** | **0.086** | **0.086** | **17.085** | **<0.001** | *** |
| SFWC:contamination | 7 | 0.025 | 0.004 | 0.711 | 0.663 |  |
| SFWC:compartment | 7 | 0.042 | 0.006 | 1.196 | 0.308 |  |
| contamination:compartment | 1 | 0.002 | 0.002 | 0.332 | 0.565 |  |
| SFWC:contamination:compartment | 7 | 0.016 | 0.002 | 0.464 | 0.859 |  |
| Residuals | 159 | 0.799 | 0.005 |  |  |  |
| *PAH-RHD Gram Negative* | *Df* | *Sum Sq* | *Mean Sq* | *F value* | *Pr(>F)* |  |
| SFWC | 7 | 0.5 | 0.072 | 0.304 | 0.951 |  |
| **contamination** | **1** | **6.23** | **6.226** | **26.486** | **<0.001** | *** |
| compartment | 1 | 0.01 | 0.01 | 0.043 | 0.836 |  |
| SFWC:contamination | 7 | 0.68 | 0.097 | 0.411 | 0.895 |  |
| SFWC:compartment | 7 | 1.34 | 0.192 | 0.816 | 0.575 |  |
| contamination:compartment | 1 | 0.24 | 0.236 | 1.005 | 0.318 |  |
| **SFWC:contamination:compartment** | **7** | **3.91** | **0.558** | **2.375** | **0.025** | * |
| Residuals | 159 | 37.38 | 0.235 |  |  |  |
| *PAH-RHD Gram Positive* | *Df* | *Sum Sq* | *Mean Sq* | *F value* | *Pr(>F)* |  |
| SFWC | 7 | 0.0624 | 0.0089 | 1.103 | 0.364 |  |
| **contamination** | **1** | **0.723** | **0.723** | **89.547** | **<0.001** | *** |
| **compartment** | **1** | **0.4619** | **0.4619** | **57.204** | **<0.001** | *** |
| SFWC:contamination | 7 | 0.0414 | 0.0059 | 0.733 | 0.644 |  |

|  |  |  |  |  |  |
| --- | --- | --- | --- | --- | --- |
| SFWC:compartment | 7 | 0.0862 | 0.0123 | 1.525 | 0.162 |
| contamination:compartment | 1 | 0.0215 | 0.0215 | 2.667 | 0.104 |
| SFWC:contamination:compartment | 7 | 0.0122 | 0.0017 | 0.215 | 0.982 |
| Residuals | 157 | 1.2677 | 0.0081 |  |  |

† Values in bold indicate significant or marginally significant effects. The number of asterisks denote the strength of the difference: \* for p values < 0.05; \*\*\* for p values <0.001 and a dot instead of an asterisk to denote nearly significant difference. Df, degrees of freedom; F, variance ratio; Pr(>F), P value.

The bacterial and archaeal Shannon H’ diversity was significantly influenced by contamination and the soil compartment (Table 3). PHE pots and rhizosphere soils were more diverse as compared to CTRL and bulk compartments. Interestingly, the rhizosphere of PHE CN (5.58), CEN (5.48), CE (5.45) and EN (5.44) had the highest Shannon H’ diversity (other values ranging from 5.41 to 4.72). However, the only two significantly different treatments in *post-hoc* tests were the rhizosphere of PHE CN treatments, compared to the bulk of CTRL C (Table 2).

The Shannon H’ diversity of Gram-negative bacterial PAH degraders experienced a significant effect of the triple interaction term (Table 3). In this sense, the bulk of PHE CEN presented the highest median Shannon H’ diversity (Table 2), followed by the rhizosphere of PHE CEN and the rhizosphere of CTRL BF.

Regarding the Gram-positive bacterial degraders, the Shannon H’ index was also significantly higher in the PHE pots and the rhizosphere soil compartment (Table 3), in line with the results for the *Bacteria* and *Archaea* general community. All the treatments with nematodes (N) had the highest absolute diversity (Table 2).

### 3.3. Microbial community structure

The fungal community structure was mainly shaped by contamination (R^2^ = 5.3%, *p* < 0.001), and the soil food web complexity (SFWC; R^2^ = 6.1%, *p* = 0.007; Figure 2A). The PCoA showed CTRL (left side) and PHE (right side) pots dispersed along the first axis. The second axis allowed for a greater dissemination of CTRL fungal community structures, compared to PHE pots (Figure 2A), along with the separation of pots containing collembolans (C and CN) from the ones containing earthworms and/or nematodes (E, N, CE and EN).

**Figure 2.**
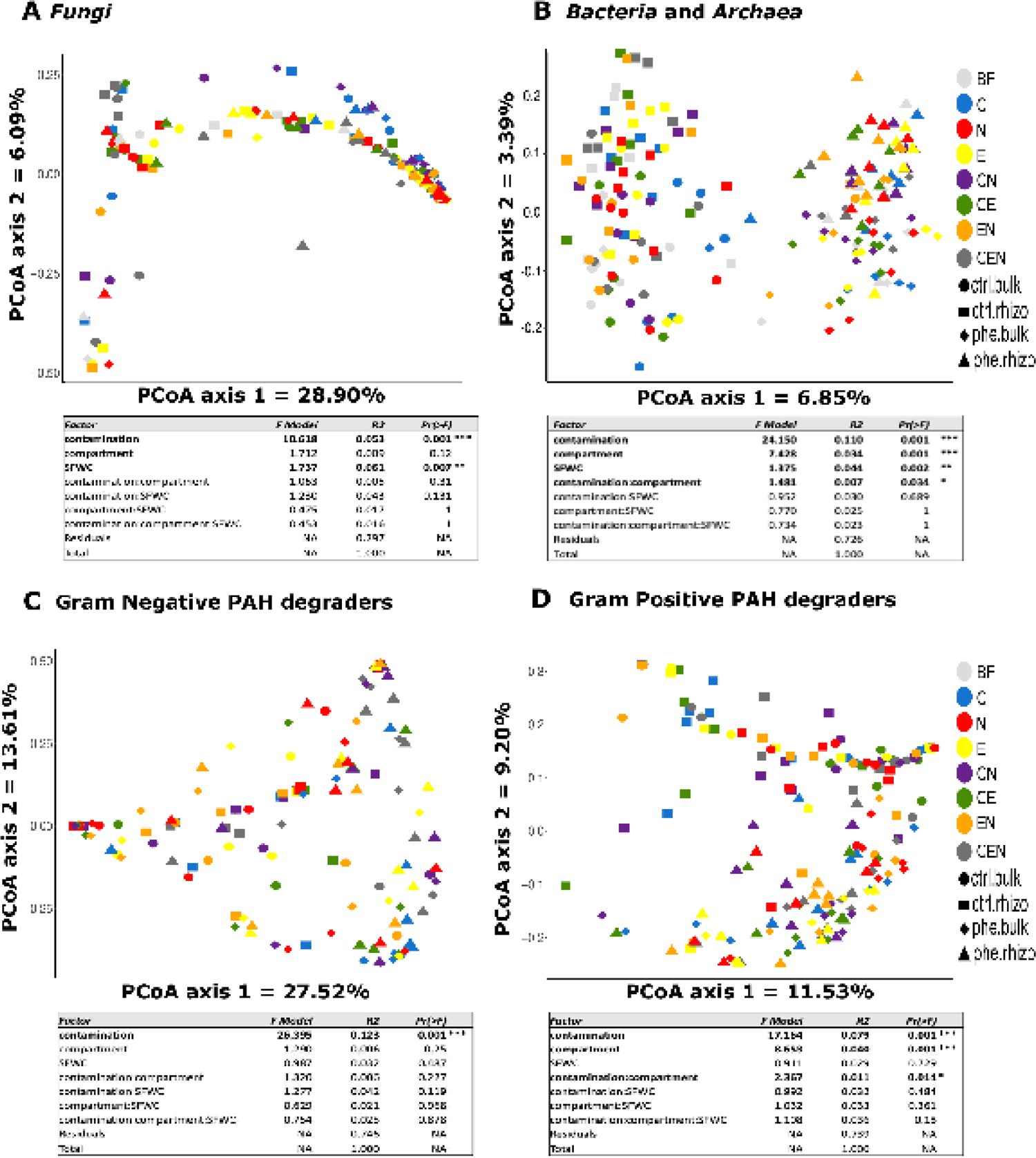
Principal coordinate analysis (PCoA) based on Bray-Curtis dissimilarity of the relative abundance of ASV showing the effects of contamination, soil food complexity treatment, soil compartment and the respective interactions of these three factors on the community structures of **A)** *Fungi* (based on the ITS region); **B)** *Bacteria* and *Archaea* (based on the 16S rRNA gene); **C)** Gram Negative bacterial degraders (based on the PAH-RHDα GN gene) and **D)** Gram Positive bacterial degraders (based on the PAH-RHDα GP gene). Below each PCoA there is the summary table of the permutational multivariate analysis of variance (PERMANOVA) examining the differences in the microbial communities based on the above-mentioned factors. Legend: SFWC levels: BF = only naturally present *Bacteria* and *Fungi*; CEN = BF plus added collembola, nematodes and earthworms; C = BF and added collembola; N = BF and added nematodes; E = BF and added earthworms; CE = BF and added collembola and earthworms; CN = BF and added collembola and nematodes; EN = BF and added nematodes and earthworms; con.bul: bulk compartment in control pot; con.rhi: rhizosphere compartment in control pot; phe.bul: bulk compartment in phenanthrene contaminated pot and phe.rhi: rhizosphere compartment in phenanthrene contaminated pot.

For *Bacteria* and *Archaea*, the PERMANOVA analysis (Figure 2B), showed that the contamination (R^2^ = 11.0%, *p* < 0.001), the soil compartment (R^2^ = 3.4%, *p* < 0.001) and the SFWC treatment (R^2^ = 4.4%, *p* = 0.002), had all significant main effects on the community structure. In addition, the effect of the contamination was modulated by the soil compartment (R^2^ = 0.7%, *p* = 0.034). Most of these effects are visible in the PCoA (Figure 2B), as the communities separate along the first axis in two clear groups, with CTRL soils on the left and PHE soil on the right. These clusters further separate between the rhizosphere and bulk soil samples along the second axis. As for the interaction between the contamination and the soil compartment, the samples within the bulk soil and the rhizosphere of the CTRL treatments were more dispersed than the samples from bulk and rhizosphere of the PHE treatment, respectively. The changes in the community structure caused by the SFWC were not evident on the first two axis of the ordination and the specific effects are presented in the community composition section of this manuscript.

The PERMANOVA analysis of the PAH-RHDα GN gene dataset showed that contamination was the only factor significantly affecting the community structure (R^2^ = 12.3%, *p* < 0.001; Figure 2C). As opposed to the ordination of the bacterial and archaeal community structures, the PAH-RHDα GN gene dataset did not cluster into clearly identifiable groups. However, it is possible to detect some separation between PHE (right) and CTRL (center-left).

Finally, the PCoA ordination of the PAH-RHDα GP gene dataset showed a stronger separation between PHE (bottom) and CTRL (top) communities (Figure 2D). Again, the main effect shaping the community structure was the contamination level (R^2^ = 7.9%, *p* < 0.001), but the compartment (R^2^ = 4.0%, *p* < 0.001) and its interaction with contamination (R^2^ = 1.1%, *p* < 0.001; Figure 2D) had also significant effects (Figure 2D).

### 3.4. Community composition

A total of 5,408 fungal ASVs were retained after sequence data processing in the bioinformatic pipeline. As for the community structure, the fungal community composition was principally influenced by contamination. The ASVs were classified in 14 phyla, with *Ascomycota* being the principal phylum. These phyla were further classified in 41 classes and 599 genera. Out of this, 14 genera had a relative abundance higher than 0.5%, and accounted in average for 67.44 to 98.83% of the total ASVs in each treatment combination. However, most pots were vastly dominated by the genus *Sphaerosporella* (Class Pezizomycetes; Figure 3A). The contamination significantly decreased the relative abundances of *Alternaria*, *Cladosporium*, *Cystobasidium*, *Gibberella*, *Mortierella*, other *Ascomycota*, other *Fungi*, other *Ostropales*, *Rhodotorula* and *Zopfiella* and only increased the relative abundance of *Sphaerosporella* (Figure 3A and Suppl. Table 3). The soil compartment affected the relative abundance of fewer genera, with the bulk soil harboring relatively more *Cystobasidium* and *Rhodotorula* whereas the relative abundance of *Mortierella*, other *Ascomycota* and other *Fungi* was higher in the rhizosphere (Figure 3A and Suppl. Table 3). Some genera were influence by the SFWC treatment. *Chaetomium*, *Sphaerosporella* and *Zopfiella* were significantly affected by the SFWC treatment, whereas *Cystobasidium*, other *Ascomycota* and other *Fungi* were marginally affected (Figure 3A and Suppl. Table 3). The relative abundance of *Chaetomium* was higher in treatments with collembolans, with mean relative abundances of 10.04%, 7.26%, 7.60% and 4.41% in C, CE, CEN and CN treatments, respectively, whereas its relative abundance in the rest of the treatments did not attain 1%. The relative abundance of *Zopfiella* increased in the presence of earthworms and collembolans, with mean relative abundances of 4.36%, 8.13%, 8.26% and 0.29% in C, CE, CEN and CN treatments, respectively. Again, in the E, EN, N and BF treatments *Zopfiella* did not present relative abundances higher than 1%. The changes in the relative abundance of *Sphaerosporella* could not be attributed to a single animal or a specific combination of animals, with the lowest mean relative abundances appearing in the CE (53.17%), N (54.08%) and CEN (56.38%) treatments and the highest in the CN (80.71%) and EN (77.57%) treatments.

**Figure 3.**
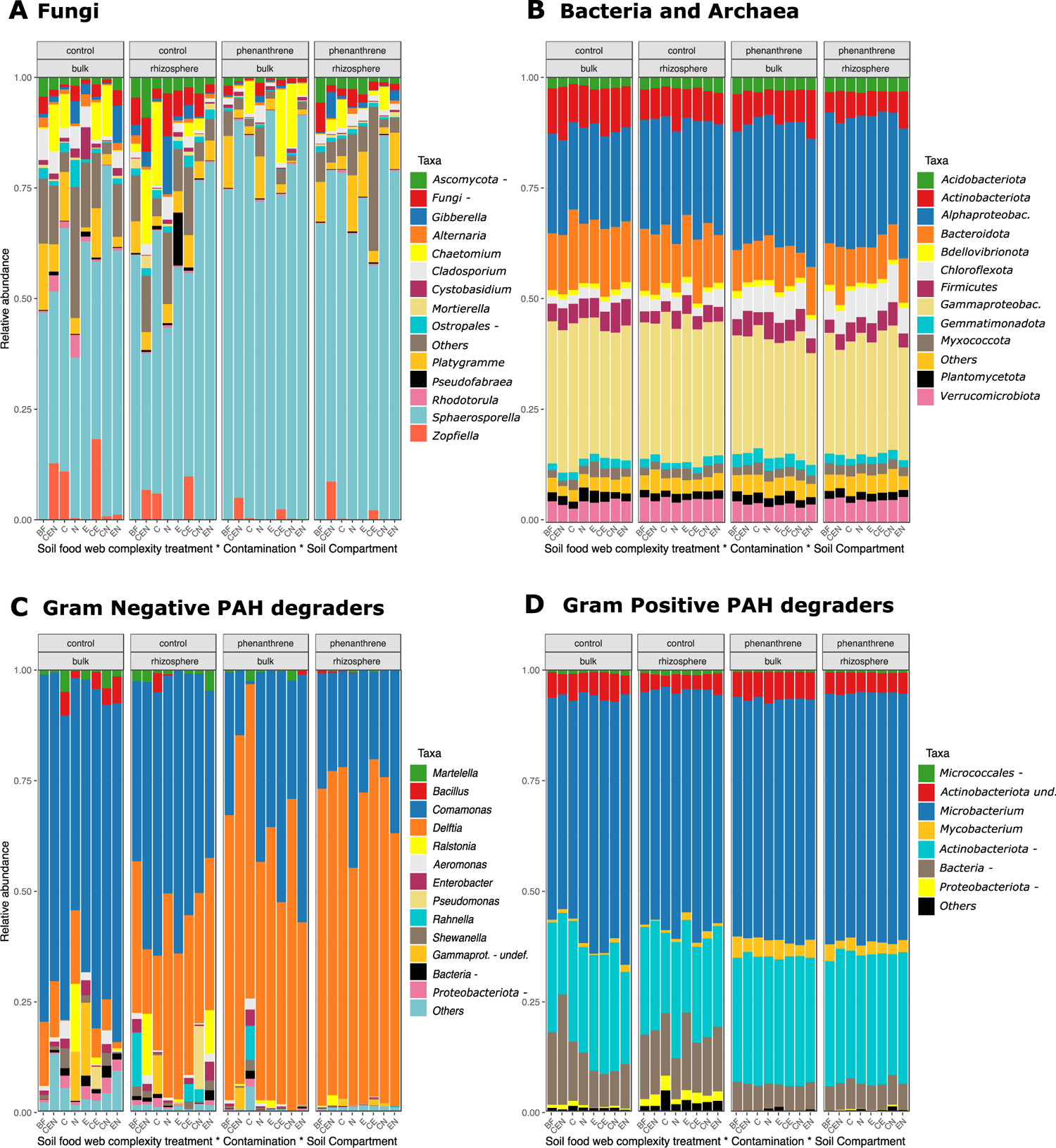
Microbial community composition based on the relative abundance of ASV. Values are averaged across treatments. Only taxa with a relative abundance above 0.1% are shown. **A)** Fungal community composition at the genus level. **B)** Bacterial and archaeal community composition at the phylum level. **C)** Gram negative bacterial degraders community composition at the genus level. **D)** Gram positive bacterial degraders community composition at the genus level. Legend: SFWC levels: BF = only naturally present *Bacteria* and *Fungi*; CEN = BF plus added collembola, nematodes and earthworms; C = BF and added collembola; N = BF and added nematodes; E = BF and added earthworms; CE = BF and added collembola and earthworms; CN = BF and added collembola and nematodes; EN = BF and added nematodes and earthworms. The “-*”* symbol represents other members of the taxa they accompany (ej. “*Ostropales-”* means “other *Ostropales”*).

A total of 15,154 ASVs were obtained from the 16S rRNA gene amplicon libraries. Those ASVs were classified in 50 phyla, where 5 belonged to *Archaea*, clustering into a total of 86 ASVs (*Crenarchaeota*, *Euryarchaeota*, *Halobacterota*, *Nanoarchaeota* et *Thermoplasmatota*). The rest of the 45 phyla belonged to *Bacteria*, but the majority of the ASVs belonged to *Acidobacteriota*, *Actinobacteriota*, *Bacteroidota*, *Bdellovibrionota*, *Chloroflexota*, *Firmicutes*, *Gemmatimonadota*, *Myxococcota* (previously classified as an order of *Deltaproteobacteria*)*, Planctomycetota, Proteobacteriota* and *Verrucomicrobiota*.

The main effects among *Bacteria* at phylum level relative abundances were caused by contamination and compartment (Suppl. Table 4 and Figure 3B). Specifically, the phyla *Actinobacteriota* (8.48% vs 6.63%), *Bacteroidota* (13.77% vs 9.18%), and *Gammaproteobacteriota* (30.94% vs. 26.47%) had significantly higher relative abundances in CTRL soils as compared to PHE soils. In contrast, *Acidobacteriota* (2.45% vs 3.15%), *Alphaproteobacteria* (22.85% vs. 28.07%), *Chloroflexota* (2.54% vs 6.14%)*, Gemmatimonadota* (1.76% vs. 2.37%) and *Mixococcota* (2.08% vs 2.25%) had higher relative abundances in PHE soils (Suppl. Table 4 and Figure 3B). There were significant changes in relative abundance of bacterial phyla due to soil compartment. For instance, *Actinobacteriota, Bacteroidota, Firmicutes*, *Gammaproteobacteria*, *Gemmatimonadota*, and *Planctomycetota* had slightly higher relative abundances in bulk soils compared to the rhizosphere. In contrast, *Chloroflexota* and *Verrucomicrobiota* had higher relative abundance in the rhizosphere. Even though the SFWC treatment did not significantly affect the composition of the general bacterial and archaeal community at the phylum level, it modulated the response of *Actinobacteriota* and *Bacteroidota* to the contamination level (Suppl. Table 4). For instance, the CEN and the EN treatments increased the relative abundances of *Actinobacteriota* and *Bacteroidota* in the rhizosphere of PHE pots up to levels similar to CTRL, while in bulk soils, the C and the CN treatments reduced the relative abundance of *Bacteroidota* in PHE pots (Suppl. Figure 1).

After quality filtering, 2,818 ASVs were retained for the PAH-RHDα GN genes, classified into 3 phyla (*Deinococcota*, *Firmicutes*, *Proteobacteriota*), 6 classes (*Alphaprotebacteria*, *Betaproteobacteria*, *Gammaproteobacteria*, *Bacilli* and *Deinococci* and other *Bacteria*) and 22 genera. From the later, 13 genera had a relative abundance higher than 0.5% (Suppl. Table 5): (*Aeromonas*, *Bacillus*, *Comamonas*, *Delftia*, *Enterobacter*, *Gammaproteobacteria*-undef, *Martelella*, *Pseudomonas*, *Rahnella*, *Ralstonia*, *Shewanella*, other *Bacteria* and other *Proteobacteria*). Contamination was the main factor determining the changes in the relative abundance of these genera, some were also affected by the soil compartment, and none were affected by the SFWC treatment (Suppl. Table 5 and Figure 3C). The phenanthrene reduced the relative abundance of *Aeromonas* (1.19% vs. 0.30%), *Bacillus* (1.39% vs. 0.17%), *Comamonas* (59.68% vs. 30.60%), *Enterobacter* (1.40% vs. 0.31%), *Martelella* (1.79% vs. 0.47%), other *Bacteria* (0.98% vs. 0.19%), other *Proteobacteriota* (1.24% vs. 0.18%) and *Shewanella* (1.32% vs. 0.28%). Only the relative abundance of *Delftia* was increased in contaminated pots both in the bulk (7.67% vs. 60.87%) and the rhizosphere (31.46% vs. 69.68%) compartments. The compartment also affected the relative abundance of both *Comamonas* and *Delftia*. The former represented on average 49.77% of the relative abundance of all bulk compartments and 39.79% of the rhizosphere soils. *Delftia*, on the other hand, represented in average 50.77% of the relative abundance in the rhizosphere compartment compared to the 36.08% of the bulk compartment.

For PAH-RHDα GP genes, a total of 6,415 ASVs were retained after quality filtering, belonging to 9 phyla (*Actinobacteriota*, *Bacteroidota*, *Chloroflexota*, *Firmicutes*, *Planctomycetota*, *Proteobacteriota*, *Spirochaetota*, *Verrucomicrobiota*), 18 classes and 67 genera. Only 7 genera had a mean relative abundance higher than 0.05% (Suppl. Table 6 and Figure 3D). *Microbacterium* was the only genus not affected by contamination, compartment or SFWC, representing approximately 53 to 56% of all PAH-RHDα GP ASVs. The contamination was again the main factor changing the mean relative abundance of the other genera. All Actinobacteria genera increased their mean relative abundance in contaminated soils compared to controls, including *Mycobacterium* (0.83% vs. 3.20%), undefined *Actinobacteria* (4.60% vs. 5.48%) and other *Actinobacteria* (23.76% vs. 29.05%). On the other hand, other *Bacteria*, (13.18% vs. 5.92%), other *Micrococcales* (0.85% vs. 0.50%) and other *Proteobacteriota* (1.30% vs. 0.10%) decreased their relative abundance in contaminated pots compared to controls. The mean relative abundance of other *Bacteria* was slightly affected by the SFWC treatment, with a higher relative abundance in the CEN (12.94%) compared to the BF (10.61%) and the other treatments (ranging from 7.36% to 9.73%).

### 3.5. PAH-RHDα GN and GP gene abundance

The absolute abundance of GP and GN genes differed by soil compartment and contamination level (Figure 4 and Table 4). The contamination with phenanthrene increased the absolute abundance of PAH-RHDα genes for both Gram-positive and Gram-negative bacteria (Table 4). However, the compartment effect was stronger in PAH-RHDα Gram-Negative genes, with the rhizosphere presenting a higher relative abundance compared to bulk soil. For PAH-RHDα Gram-Positive genes, the compartment effect was weaker, with a slightly higher relative abundance in the bulk soil compared to the rhizosphere (Figure 4 and Table 4). There was no significant effect of the SFWC treatment (Table 4).

**Figure 4.**
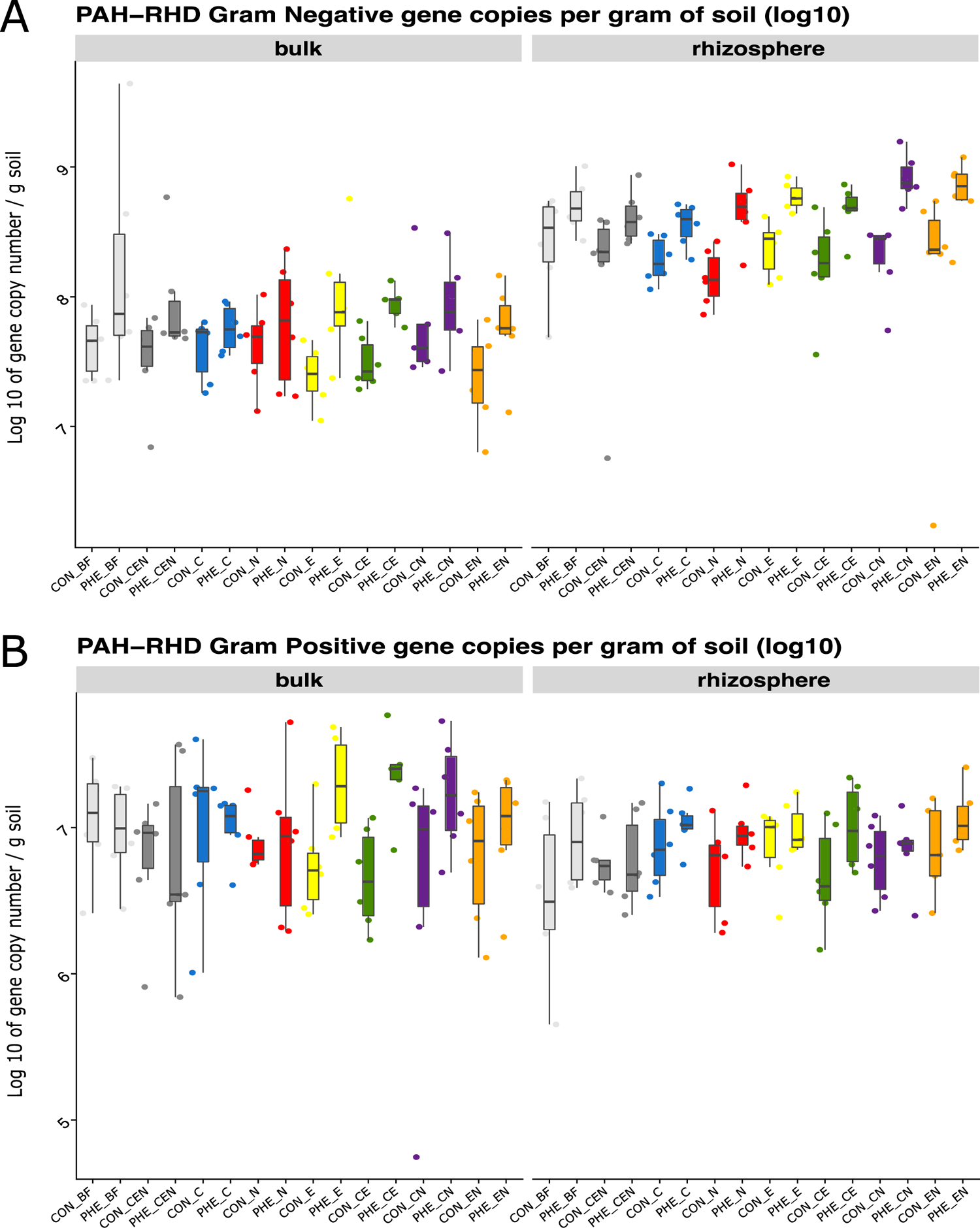
PAH-RHDα gene copy numbers determined by real-time PCR quantification on DNA for **A)** Gram negative bacterial degraders and **B)** Gram positive bacterial degraders. Values are log transformed. Legend: CON_*: control pots; PHE_*: phenanthrene contaminated pots; SFWC levels: BF = only naturally present *Bacteria* and *Fungi*; CEN = BF plus added collembola, nematodes and earthworms; C = BF and added collembola; N = BF and added nematodes; E = BF and added earthworms; CE = BF and added collembola and earthworms; CN = BF and added collembola and nematodes; EN = BF and added nematodes and earthworms.

**Table 4.**
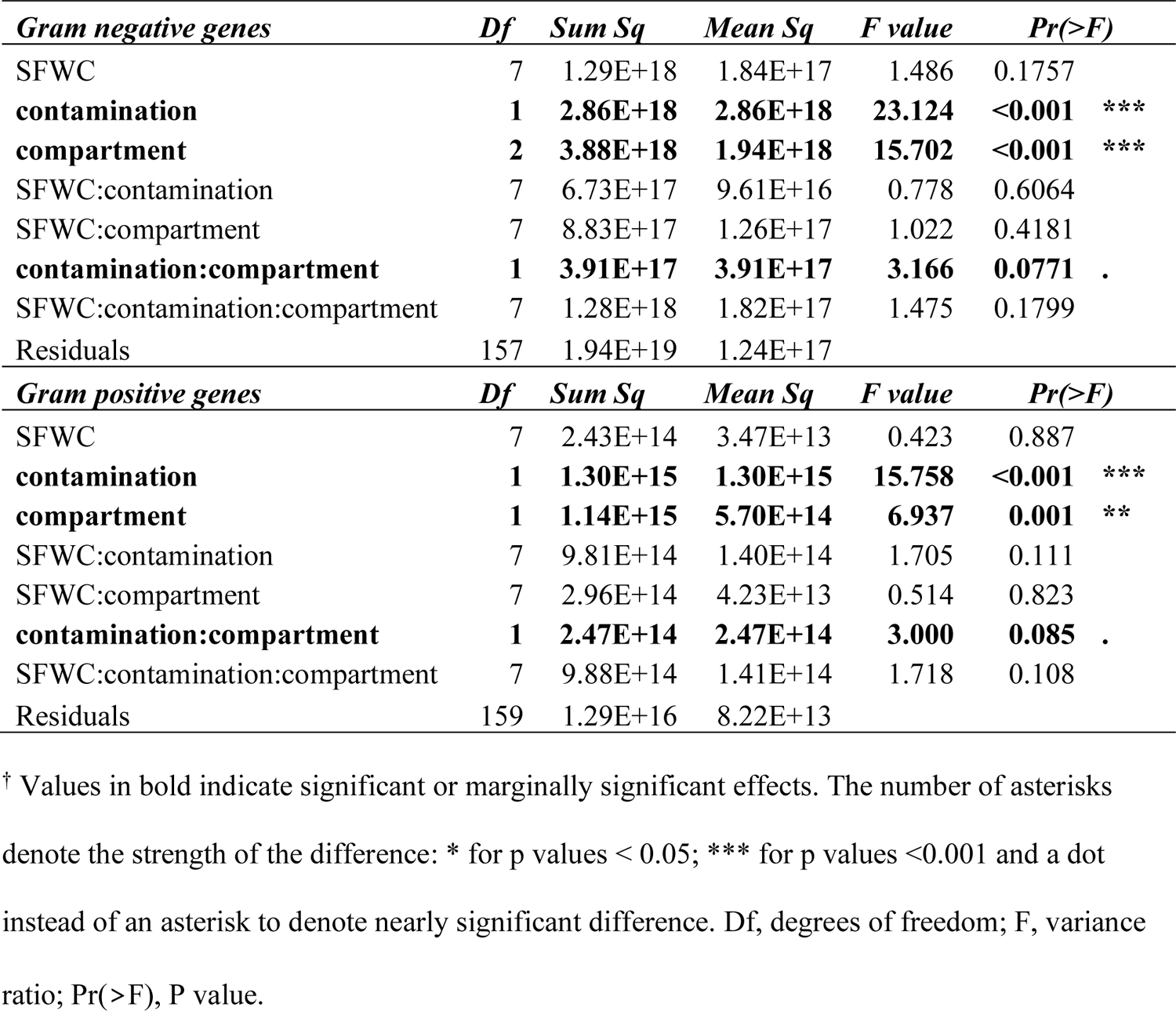
Summary of the three-way analysis of the variance (ANOVA) on the absolute abundance of PAH-RHD genes (on copy numbers per gram of soil) ^†^.

## 4. Discussion

Although there is the ample evidence that the soil fauna strongly affects the soil microbial communities (Bonkowski et al., 2009; Cheng et al., 2011; Jing et al., 2017; Yeates, 2007) and that these latter are the central actors during contaminant degradation (Correa-García et al., 2018), little evidence is available linking soil fauna to phytoremediation efficiency and the underlying microbial communities. In this study, we aimed to determine the effects of adding collembolans, nematodes and earthworms to soils on the phytoremediation of phenanthrene by willows.

Our first hypothesis was that the willows subjected to the highest level of soil food web complexity (SFWC) treatment would be more productive than the willows exposed to lower levels of SFWC. Our results related to this hypothesis are not conclusive. Indeed, all faunal treatments resulted in taller trees that produced more biomass as compared to the microbes-only contaminated pots, but because of the high variability between replicates, this was mostly not statistically significant. Significant differences in biomass and length were observed between the willows growing in contaminated soils in the presence of microbes only and the willows growing in the non-contaminated soils and inoculated with either collembolans, nematodes and earthworms or earthworms and nematodes. This is probably the result of a combination of a decreased growth due to phenanthrene contamination and an increased growth due to the inoculation of soil fauna. Also, gamma radiation may have significantly reduced the initial microbial community hence contributing to a decrease in willow growth. But since no microbial analyses were performed before or after irradiation, this explanation remains hypothetical.

Košnář, Mercl and Tlustoš (2020) found significantly lower biomass production for willows (*Salix smithiana*) growing in PAHs contaminated pots compared to controls. A field study looking at higher concentrations of mixed contaminants also showed significant decreases of the willow biomass in contaminated plots compared to controls (Brereton et al., 2016). Our results are also in line with the reported positive effects of earthworms on plant productivity (Blume-Werry et al., 2020; Van Groenigen et al., 2014), including during phytoremediation of lead (Jusselme et al., 2015). This effect has been previously attributed to the bioturbation (burrowing) activity of earthworms, that results in many beneficial effects for plants, such as improved soil porosity (Shipitalo and Le Bayon, 2004) leading to increased soil water infiltration (Bouché and Al-Addan, 1997) and aeration (Bartlett et al., 2010), organic matter degradation and increased nutrient cycling (Edwards, 1997). Bacterivorous nematodes, such as the one used in this study, also have the potential to stimulate plant growth through an increased mineralization of soil nutrient (Alphei et al., 1996).

Our second hypothesis was that the microbial communities would be affected by the SFWC treatments. We confirmed this hypothesis as the SFWC had a relatively small but significant effect on the community structure and composition of Bacteria and Archaea, but not on its diversity indexes. Previous studies have shown that remediation rates and plant growth can be linked to the initial soil microbial community composition and structure (Bell et al., 2016, 2015; Correa-García et al., 2021; Yergeau et al., 2015). The shifts in the bacterial and archaeal communities caused by the SFWC treatments could therefore indirectly explain part of the trends observed for plant growth. However, this shift in microbial communities caused by the SFWC treatments did not seem to translate into different degradation rates, most probably because the PAH-degraders subset of the community was, in general, not affected by the SFWC treatments. Indeed, for PAH degraders, the only significant effect of SFWC was on the diversity of Gram-negative degraders, and on the relative abundance of a few phyla for Gram-positive degraders.

Their general community composition and structure were unaffected by SFWC, in contrast to the trends observed for Bacteria and Archaea. The abundance of both Gram-negative and Gram-positive was mostly determined by the contamination levels, in agreement with previous work (Cébron et al., 2008; Sawulski et al., 2014).

We also observed that SFWC appeared to modulate the effect of the contamination on Bacteria and Archaea (marginally significant SFWC x Contamination effects on relative abundance of some phyla). For instance, for the pots treated with earthworms and nematodes (EN) or collembolans, earthworms and nematodes (CEN) the relative abundances of the phyla *Actinobacteriota* and *Bacteroidota* were higher in the contaminated rhizosphere as compared to the control rhizosphere, effectively reversing the trend observed for all other *treatments*. These results are in agreement with the increased relative abundance reported for Bacteroidota in the presence of the earthworms *Pontoscolex corethrurus* (Bernard *et al*. 2012), *Eisenia fetida* (Cai et al., 2018), *Lumbricus terrestris* and *Aporrectodea caliginosa* (Nechitaylo et al., 2010). An increased relative abundance of *Actinobacteriota* in soils after their passage through the earthworm gut has also been reported (Aira et al., 2016; Cai et al., 2018; Medina-Sauza et al., 2019; Sun et al., 2020). The trends observed for *Actinobacteriota* were also visible in the significant effect of SFWC (in interaction with other factors) on Gram-positive PAH degraders related to this phylum. This specific effect of SFWC on *Actinobacteriota* could partially explain why earthworms have proven to be especially useful in PAH remediation (Contreras-Ramos et al., 2006; Coutiño-González et al., 2010; Hernández-Castellanos et al., 2013).

As for *Fungi*, we found a significant effect of the SFWC treatments on the community structure, composition, and diversity. Similarly, the presence of earthworms have been previously related to higher fungal diversity during lead phytoremediation (Jusselme et al., 2015). In our study, the shifts caused by the SFWC treatments seemed to be mainly driven by *Chaetomium, Zopfiella* and *Sphaerosporella*. For instance, *Chaetomium* was found to be significantly more abundant in the presence of collembolans. Accordingly, several springtail species (*Ceratophysella denticulata*, *Folsomia candida*, *Protaphorura fimata*, *Sinella curviseta*) have been found to graze on *Chaetomium globosum* hyphae and to participate to the dispersion of their spores (Erktan et al., 2020b; Haque, 2018; Haubert et al., 2008). *Chaetomium* had also been reported as predominant in greenhouse mesocosms aiming to degrade oxytetracycline with earthworms and arbuscular mycorrhizal fungi (Cao et al., 2015). Also, in our experiment, the genus *Zopfiella* was relatively more present in treatments containing a combination of collembolans and earthworms. This is in line with previous reports that found that *Zopfiella* was particularly present during vermicomposting (Neher et al., 2013). In contrast, *Zopfiella* was found to account for up to 5% of the relative abundance of fungi in control farmland soils, which was significantly higher than in the same soils treated with earthworms (Song et al., 2020).

Some *Zopfiella* species produce zopfiellin, a secondary metabolite with antifungal properties (Futagawa et al., 2002), that could be involved in soil pathogen suppression (Liu et al., 2019) in some soils, and might have indirectly affected the fungal community in our experiment, explaining indirectly some of the SFWC effect on fungal communities. Furthermore, collembolans have been found to feed preferably on pathogenic fungi rather than on mycorrhizal hyphae (Innocenti and Sabatini, 2018). In our study, a similar behavior could explain the higher relative abundance of *Sphaerosporella* in more complex SFWC treatments.

Finally, we hypothesized that willows growing in soils with a higher SFWC would rhizodegrade more phenanthrene than willows growing in soils with a lower SFWC. Although we did find some effect of SFWC on microbial communities and willow growth, which are the two most important factors for rhizoremediation efficiency, our results disproved this hypothesis as there were no significant differences due to the SFWC treatments in the amount of phenanthrene found in soils after 8 weeks of willow growth. As mentioned above, this could be due to the general lack of effect of the SFWC treatments on the PAH-degraders. These results are in sharp contradiction with previous studies that reported a positive direct effect of soil fauna, especially earthworms, on the degradation of different types of pollutants (Contreras-Ramos et al., 2006; Coutiño-González et al., 2010; Hernández-Castellanos et al., 2013). A recent study from our group showed that initial soil physicochemical and microbiological characteristics were critical for effective phytoremediation, with poplar trees only significantly affecting phenanthrene degradation in one of the two soils tested (Correa-García et al., 2021). Similarly, it could be well possible that because of the characteristics of the soil used here, the effect of SFWC on degradation rates has gone unnoticed. In fact, in most pots, both for rhizosphere and bulk soil, over 95% of the applied phenanthrene was degraded, suggesting that plant presence was not effective in enhancing degradation under our conditions. Alternatively, the unique sampling point after eight weeks of growth might have precluded the observation of any differences in degradation that might have occurred before that. Indeed, the addition of root exudates to phenanthrene contaminated sand resulted in significant reductions of the phenanthrene contamination compared to the unamended controls after only 10 days (Louvel et al., 2011). Other studies have shown a complete degradation of phenanthrene by willows within 3 months (Hultgren et al., 2010), 2 months (Correa-García et al., 2021) or as little as 21 days (Tiralerdpanich et al., 2018). Additional studies using different SFWC levels where the contaminant is evaluated at several time points throughout the experiment and in soils with different biological and physico-chemical characteristics will be needed to rule out if soil fauna influences the degradation rates during phytoremediation.

Even though most of the contaminant had disappeared, large differences in microbial communities, including PAH degraders abundance, diversity and community composition, were still visible between contaminated and control pots, suggesting some level of legacy effect of contamination. The phenanthrene contamination triggered a general reduction of the fungal diversity, which was driven by large increases in the relative abundance of *Sphaerosporella*. The predominance of this fungus has already been reported in other pot and field phytoremediation experiments carried out with trees (Bell et al., 2014a; Correa-García et al., 2021; Dagher et al., 2020; Danielson, 2007; Yergeau et al., 2015). Furthermore, its presence in the rhizosphere or in the bulk soil of pots with tree cuttings has been related to an enhancement of plant biomass (Yergeau et al., 2015) and to resistance to contamination stress (Bell et al., 2014a), probably through its role as an ectomycorrhiza (Danielson, 2007). In our study, *Sphaerosporella* relative abundance was significantly higher in contaminated pots, supporting the premise that this fungus may be helping *Salicaceae* trees survive in stressful environments. Quite interestingly, *Sphaerosporella* relative abundance was also significantly affected by the SFWC treatments, being generally higher in more complex treatments, which could contribute to explain the effects of SFWC on plant growth parameters.

## 5. Conclusion

To date, most research on phytoremediation of polycyclic aromatic hydrocarbons has focused on testing the response and the effect of soil microbial communities and the role of the plant in phytoremediation processes, yielding inconclusive results (Guo et al., 2018; Kuppusamy et al., 2017). In fact, the reality is that soils are very heterogeneous ecosystems harbouring rich communities. Meso- and macroorganisms can significantly alter the soil physicochemical environment as well as the microbial communities (Bartlett et al., 2010; Crowther et al., 2011; Erktan et al., 2020b; Ngosong et al., 2014; Yang and van Elsas, 2018). Importantly, trophic interactions are one of the mechanisms responsible for shifts in microbial communities with possible implications for phytoremediation (Li et al., 2015; Zeb et al., 2020). Our study is a first step toward an increased understanding of these trophic interactions, as our unique experimental design allowed us to disentangle the effects of the various components of the soil food web on phytoremediation efficiency. Despite the large effect of soil food web complexity (SFWC) on microbial communities, we did not find strong community shifts among PAH degraders.

Similarly, phenanthrene degradation was unaffected by SFWC treatments. Instead, we found that some levels of the SFWC treatment, namely those including earthworms, increased the willow growth, as others have reported (Rajapaksha et al., 2014). However, we did not find that bigger plants were correlated with higher degradation rates, in contrast with other studies (Dagher et al., 2020; Sun et al., 2013).

Our results confirmed that certain microbial taxa related to bioremediation success can be affected by the presence of complex soil food webs. Particularly, we found that complex levels of SFWC increased the relative abundance of the fungi *Sphaerosporella* (Bell et al., 2015; Dagher et al., 2020)*, Chaetomium* (Cao et al., 2015) and *Zopfiella* (Liu et al., 2019), as well as the abundance of actinobacterial PAH degraders (Thomas et al., 2019). These microbial community changes are not trivial and suggest potential ways by which SFWC could affect phytoremediation outcomes. We propose that the soil fauna contributes indirectly to the success of phytoremediation and should be considered in future experiments to understand the mechanisms behind their effect on soil microbial communities.

## 7. Acknowledgments

We thank the precious help provided by Denis Lachance during the setup of the experiment, as well as all the members of the Laurentian Forestry Centre and the Centre Armand Frappier Santé Biotechnologie who contributed with ideas and discussions to improve the outcome of this study. We also thank the logistic support provided by Stepan Pasharyan during the experimental set up.

## 8. Authors contributions

S.C.G., Conceptualization, Methodology, Experimental Setup, Field Work, Data collection, Data curation, Data Analysis, Writing–Original draft preparation. V.C., in-house PHE extraction methodology, Field Work, Data Collection, Original draft edition. J.A.D and E.M.K, Experimental Setup, Field Work, Data Collection, Original draft edition. J.T., Bioinformatic Analysis and Bioinformatic methods writing, Original draft edition. A.S., Conceptualization, Methodology, Supervision, Reviewing, and Editing. E.Y., Funding, Conceptualization, Methodology, Supervision, Writing, Reviewing, and Editing.

